# Variable freshwater influences on the abundance of *Vibrio vulnificus* in a tropical urban estuary

**DOI:** 10.1101/2021.09.22.461452

**Authors:** Olivia D. Nigro, La’Toya I. James-Davis, Eric Heinen De Carlo, Yuan-Hui Li, Grieg F. Steward

## Abstract

To better understand the controls on the opportunistic human pathogen *Vibrio vulnificus* in warm tropical waters, we conducted a year-long investigation in the Ala Wai Canal, a channelized estuary in Honolulu, HI. The abundance of *V. vulnificus* as determined by qPCR of the hemolysin gene (*vvhA*), varied spatially and temporally over four orders of magnitude (≤ 3 to 14,000 mL^-1^). Unlike in temperate and subtropical systems, temperatures were persistently warm (19–31°C) and explained little of the variability in *V. vulnificus* abundance. Salinity (1–36 ppt) had a significant, but non-linear, relationship with *V. vulnificus* abundance with highest abundances (> 2,500 mL^-1^) observed only at salinities from 7 to 22 ppt. *V. vulnificus* abundances were lower on average in the summer dry season when waters were warmer but more saline. Highest canal-wide average abundances were observed during a time of modest rainfall when moderate salinities and elevated concentrations of reduced nitrogen species and silica suggested a groundwater influence. Distinguishing the abundances of two genotypes of *V. vulnificus* (C-type and E-type) suggest that C-type strains, which are responsible for most human infections, were usually less abundant (25% on average), but their relative contribution was greater at higher salinities, suggesting a broader salinity tolerance. Generalized regression models suggested up to 67% of sample-to-sample variation in log-transformed *V. vulnificus* abundance was explained (n = 202) using the measured environmental variables, and up to 97% of the monthly variation in canal-wide average concentrations (n = 13) was explained with the best subset of four variables.

**IMPORTANCE:** Our data illustrate that, in the absence of strong seasonal variation in water temperature in the tropics, variation in salinity driven by rainfall becomes a primary controlling variable on *V. vulnificus* abundance. There is thus a tendency for a rainfall-driven seasonal cycle in *V. vulnificus* abundance that is inverted from the temperature-driven seasonal cycle at higher latitudes. However, stochasticity in rainfall and its non-linear, indirect effects on *V. vulnificus* concentration means that high abundances can occur at any location in the canal at any time of year, making it challenging to predict concentrations of this pathogen at high temporal or spatial resolution. Much of the variability in canal-wide average concentrations, on the other hand, was explained by a few variables that reflect the magnitude of freshwater input to the system, suggesting that relative risk of exposure to this pathogen could be predicted for the system as a whole. [at 148 out of 150 words max]

## INTRODUCTION

The bacterium *V. vulnificus* is an opportunistic and formidable human pathogen that has a world-wide distribution, in a variety of marine and estuarine environments (1). In humans, *V. vulnificus* may cause a range of illnesses that includes gastroenteritis, necrotizing fasciitis and septicemia (2). Infections occur as a result of ingestion of contaminated seafood (3) or via wound exposure to waters (4). Strains vary in their propensity to cause disease in humans, with certain genotypically distinguishable strains much more commonly, but not exclusively, associated with disease in humans (5). The exact mechanisms of virulence in *V. vulnificus*, and the genes responsible for the onset of illness, have yet to be determined, but a number of correlative biomarkers have been used to discriminate those strains most commonly associated with human disease (6). Variations in the 16S rRNA gene, for example, have been used in PCR assays to discriminate “A-type” strains from “B-type” strains (7, 8), the latter of which predominate among clinical isolates. Another commonly used marker is the 200 bp segment of the virulence-correlated gene that resolves the gene variants *vcgC* or “C-type” strains from *vcgE* or “E-type” strains (9). PCR-based analysis of fifty-five *V. vulnificus* isolates indicated that 90% of the strains isolated from infected patients were of the C-type (clinical), while 93% of the strains isolated from the environmental samples were E-type (environmental). Subsequent analyses revealed broader genomic differences along with physiological differences between these lineages suggesting that they are distinct ecotypes that may be better adapted for either environmental growth (E-type) vs. stress tolerance (C-type) (10). These biomarkers are largely congruent, with the common environmental strains being A-type/E-type, and the majority of clinical isolates being B-type/C-type, although all types can cause disease in humans (11).

Studies of *V. vulnificus* in temperate and subtropical waters have shown that warmer temperatures increase the frequency of detection (12–15). Quantification over an annual cycle reveals a clear temperature-driven seasonal signal, with the highest concentrations of *V. vulnificus* occurring in warm summer months (16–19) and culturable cells declining dramatically at temperatures below 13 °C (6). *V. vulnificus* abundance is also influenced by salinity (19–21), thriving in conditions of both warm temperatures and moderate salinities (5). The environmental patterns of abundance are consistent with observations of *V. vulnificus* growth under controlled laboratory conditions (12, 22) that show increasing growth rates up to around 37 °C, and a broad salinity tolerance with fastest growth rates between 5–25 ppt. Correlation models of environmental data support the idea that temperature and salinity are two of the most important variables controlling *V. vulnificus* abundance, but their relative importance depends on the ranges over which they are sampled. (21, 23–26).

In temperate environments, the incidence of *V. vulnificus* infection tracks the seasonal environmental abundances of the pathogen, with the most infections occurring during the warm summer months (6). It follows that the inhabitants of sub-tropical, and especially tropical areas, where air and water temperatures are warm year-round, would be particularly vulnerable to *V. vulnificus* infection. Indeed, according to available surveillance data for the years 2003–2008 (27–29), Hawaiʻ i had the fifth highest incidence of non-food-borne *V. vulnificus* infections in the U.S., trailing only four gulf states (Florida, Louisiana, Mississippi, and Texas). On a per capita basis, it was the highest in the nation. Despite higher incidence of *V. vulnificus* wound infections, primarily from recreational waters, there has been little data collected on *Vibrio vulnificus* in the coastal waters of Hawaiʻ i (30, 31) and scant data on the ecology of *V. vulnificus* in tropical waters in general (20). Consequently, we initiated an investigation of the abundance and dynamics of *V. vulnificus* in the Ala Wai Canal and Harbor.

The Ala Wai canal provides partially channelized drainage for two watersheds. Although it is not designated as a recreational waterway, the canal is used extensively for boating and fishing. Flow down the canal varies as a function of tide and of rainfall, the latter driving surface runoff (streams and storm drains) and, with some hysteresis, groundwater seepage. Salinity varies widely in the canal, as a function of depth, overall stream flow, position in the canal relative to the freshwater sources, and tidal forcing. Water temperature, on the other hand, varies over a relatively narrow range compared to temperate systems. Because of the seasonality in rainfall in Hawaii, with higher precipitation in winter months (32), we hypothesized that there could be an inverse seasonal pattern in *V. vulnificus* abundance driven by salinity compared to the strongly temperature-driven patterns in temperate waters.

Our objectives with this study were to document the temporal spatial variability of *V. vulnificus* total abundance and strain composition (C-type vs. E-Type) in the estuarine waters of the Ala Wai Canal and Harbor and to determine how abundance was related to environmental variables. The goal was to better understand the environmental controls on *V. vulnificus* in tropical estuarine waters and to assess the prospects for modeling pathogen abundance.

## MATERIALS AND METHODS

### Study Site

Sampling took place in the Ala Wai Canal (Fig. 1), a 3.1 km long, engineered waterway located on the southern coast of Oʻ ahu that separates Waikiki and urban Honolulu (33). A watershed that covers 42.4 km^2^ drains into the Ala Wai Canal via the Mānoa and Palolo Streams, which merge to form the Mānoa-Palolo Stream prior to entering the canal, and the Makiki Stream, all of which run through urban areas before reaching the canal. As a consequence, the streams are contaminated with a variety of anthropogenic substances and their convergence in the Ala Wai Canal has contributed to its pollution and eutrophication (34, 35). The influx of fresh water from the streams creates a salinity gradient with a typical salt-wedge structure. Tidal flow causes seawater to flow landward on the flood tide and seaward on the ebb tide and remain at depth. The freshwater streams flow seaward on all tides creating a freshened water surface layer estimated to extend to 0.5 m depth on average, but which is highly variable both in salinity and thickness (36). Sediments are continually deposited in the canal at the mouth of the Mānoa-Palolo Stream causing the build-up of a sill that restricts flushing of deep water in the uppermost section of the canal.

**Fig. 1.**
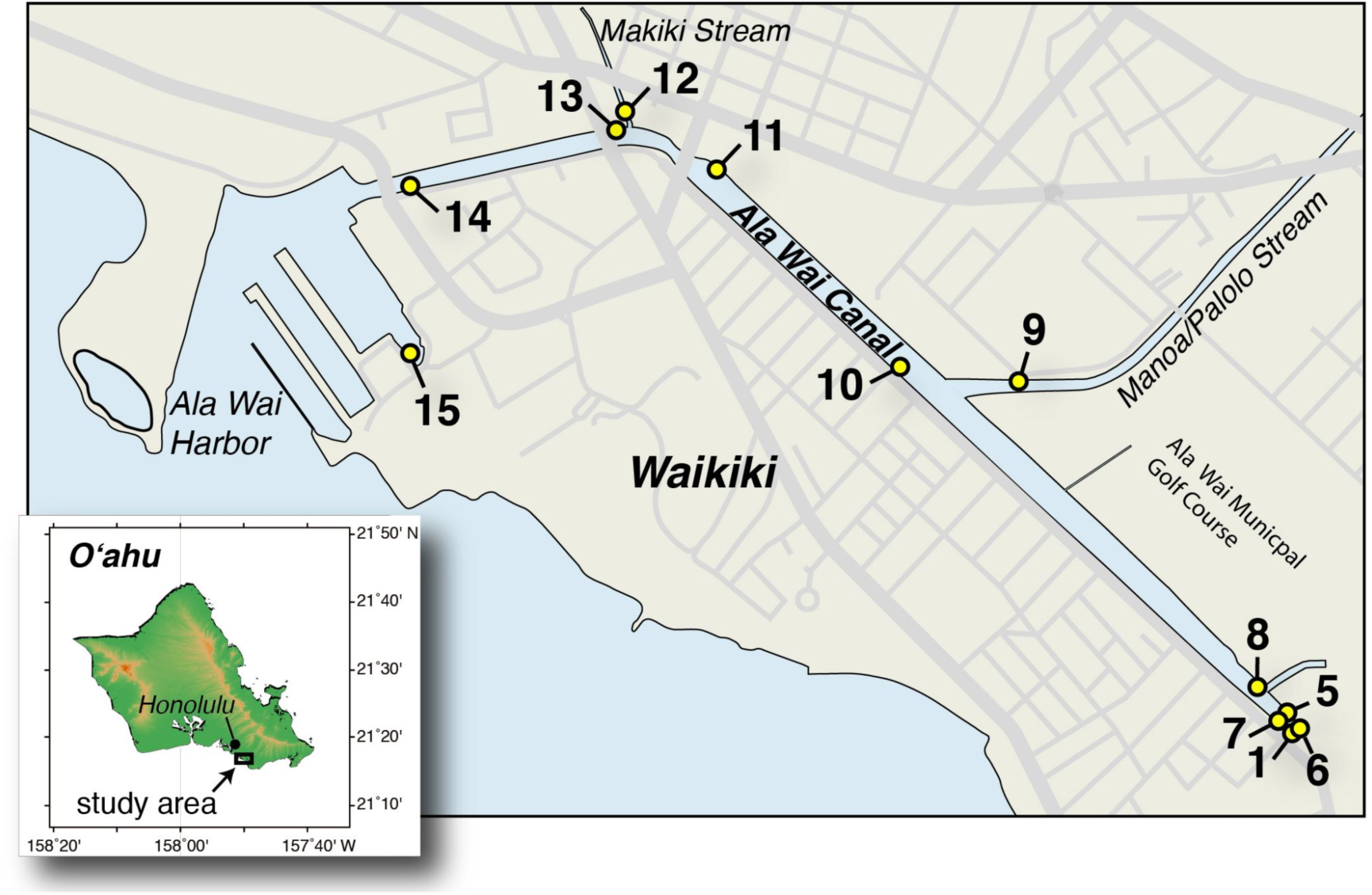
Map of Samping Sites. Inset shows the general location of the canal on the south shore of the island of Oʻ ahu in the Hawaiian Island chain. Main map shows the site numbers and position along the canal. Site 1 is at the closed end of the canal with occasional input from surface runoff via storm drains. Sites 9 and 12 are at the mouths of the Mānoa-Palolo and Makiki Streams, respectively.

### Sampling locations, dates, and times

Sampling of the Ala Wai Canal spanned 13 months beginning March 17, 2008 and concluding on March 10, 2009, covering the nominal dry summer (April– September) and rainy winter (October–March) months. Samples were collected monthly at twelve sites in the Ala Wai Canal numbered (1 and 5–15) by distance from the shallow, upper section of the canal (Site 1) to the Ala Wai Harbor (Site 15). Site 9 was just inside the mouth of Mānoa-Palolo Stream and Site 12 was at the mouth of Makiki Stream (Fig. 1, Supplemental Table S1). Missing site numbers 2–4 referred to other samplings at Site 1 that were not used in this study. Sampling at a higher temporal resolution was also conducted in the dry and rainy seasons to assess changes on shorter time scales. Samples were collected weekly at all sites for four weeks from June 26–July 17, 2008 and again for three weeks from February 22–March 10, 2009. Samples were also collected at a reduced number of sites (Sites 5, 9, 12, 14) daily for six days from July 10–15, 2008 and daily for five days from March 2–6, 2009, and once every three hours (trihoral) for twenty-four hours at Sites 5, 9, and 14 from July 15 to July 16, 2008.

### Rainfall and streamflow

Rainfall data collected by National Weather Service rain gauges (part of the Hawaiʻ i Hydronet System) at 15-minute intervals were retrieved from the online resource (https://www.weather.gov/hfo/hydronet-data). Data from two gauges were selected for analysis. The first was HI-18 (NOAA# MNLH1), which is located near the origin of Mānoa Stream (N21.3161 W157.8142) at an elevation of 150 m in Manoa Valley (“Valley” rainfall). The second is HI-26 (ALOH1), which is located at Aloha Tower (N21.3060 W157.8662) in downtown Honolulu near sea level (15 m) at the coast (“Coastal” rainfall). From these data, average daily rainfall for all sampling months was determined, as well as total rainfall from each 24-hour period prior to sampling. Data on tidal flux were obtained from the National Ocean Service (NOS), using tide gauge number 1612340. Stream flow data were obtained from the United States Geological survey (waterdata.usgs.gov/usa/nwis/uv?16247100) for the Mānoa-Palolo Stream gauge #16247100.

### Water sample collection and processing

Whole water samples were collected from the top 10–30 cm at all sites in acid-washed bottles with a pole sampler and stored on ice (except for samples used for culturing, which were kept at ~ 15 °C with cold packs) and transported to the laboratory within three hours of collection. Subsamples (ca. 25 mL) for nutrient analysis (n = 207–211) were frozen and shipped on dry ice to the Oregon State University nutrient analysis facility for determination of dissolved silica, phosphate, nitrate plus nitrite, nitrite, and ammonium concentrations (37). Nutrient concentrations were measured during every sampling event excluding two weekly sampling events in July 2008 (July 3 and 7). The values for the mean, number of samples, median, minimum and maximum of the measured nutrients have been previously reported (38).

For particulate carbon (PC) or nitrogen (PN) and chlorophyll a (chl a) measurements, subsamples (25–200 ml) were filtered onto pre-combusted glass-fiber filters (GF/F, Whatman) in duplicate and stored frozen until analysis. For PC and PN (n = 199), filters were pelletized and combusted in a high-temperature combustion CN analyzer, the CE-440 CHN elemental analyzer (Exeter Analytical) following HOT program protocols (39). Filters for chl a analysis (n = 194) were extracted in 100% acetone at -20°C for 7 days. Fluorescence of extracts and standards were measured using a Turner AU10 fluorometer before and after acidification (40).

Samples for bacteria counts (n = 219) were fixed with filtered (0.2 µm) formaldehyde (10% w/v final concentration) in a cryovial (Nalgene) and stored at - 80 °C. Total bacteria were counted by thawing samples, staining with SYBR Green I, and analyzing on an acoustic focusing flow cytometer (Attune; Thermo Fisher Scientific).

Samples for molecular analysis (100– 550 mL) were pressure filtered via peristaltic pump through 0.22 µm polyethersulfone filter capsule (Sterivex, Millipore), then stored at -80 °C until extracted.

### Cultivation on vibrio selective medium

For five of the monthly samplings (Mar, Jun, Sep, Dec 2008, and Mar 2009), water samples were filtered through 0.45 µm pore size, mixed cellulose ester filters (47 mm, GN-6; Pall) and filters were placed face-up on the vibrio-selective medium CHROMagar Vibrio (DRG Intl.). After overnight incubation, blue colonies were enumerated as putative *V. vulnificus*.

### DNA extraction and purification

DNA was extracted from the Sterivex filters using the Masterpure™ Nucleic Acid Extraction Kit (Epicentre). Six-hundred microliters of Masterpure™ Tissue and Cell Lysis Solution containing recommended quantities of proteinase K were added to each Sterivex filter. The ends of the filters were sealed and the filters incubated on a rotisserie in a hybridization oven at 65 °C for 15 minutes. Fluid was recovered from filter housing by aspiration with a syringe. The filling with buffer, incubation, and buffer recovery steps were repeated twice more and the combined extract from all three rounds was pooled (total volume ca. 1.8 ml). Three-hundred microliters of the pooled extract was processed according to the Masterpure™ Kit guidelines and the remainder was archived. Accounting for all of the raw extract volume, total DNA yields ranged from 1–540 µg L^-1^ of canal water (geometric mean of 30 µg L^-1^). Following initial purification, the resuspended DNA (200 µL) was passed through a spin column containing acid-washed polyvinylpolypyrrolidone (PVPP) in an effort to remove any residual inhibitors (41). DNA concentration in each sample was quantified fluorometrically (Quant-iT Broad Range DNA kit, Life Technologies) both before and after the PVPP purification step to account for losses incurred during the purification stage (average recovery 60%). Geometric mean concentration of DNA in the final purified extracts was 7 ng µL^-1^ (range 0.1–54 ng µL^-1^).

### Quantitative PCR

Total *V. vulnificus* was estimated by TaqMan qPCR targeting the hemolysin gene (*vvhA*) using primer and probe sequences reported by Campbell and Wright (42). Quantification of C-type *V. vulnificus* used the primers and probes targeting the virulence-correlated gene variant (*vcgC*) from Baker-Austin et al. (43). E-type *V. vulnificus* was calculated as the difference in concentration between the two assays. Both assays were prepared as 25-µL reactions with 12.5 µL of TaqMan Universal PCR Master Mix (Applied Biosystems), 1.5 µg µl^-1^ final concentration of non-acetylated bovine serum albumin (Applied Biosystems) and 0.25–0.9 µM each of the appropriate primers and probe (Supplemental Table S2), 2–5 µl of DNA template, and water as needed. For *vvhA* assay, primers were added at 0.9 µM each and the probe at 0.25 µM. For the *vcgC* assay, primers and probe were each added at 0.5 µM final concentrations. Cycling conditions consisted of initial denaturation at 95 °C (10 min), then 40 cycles of 95 °C (15 s) and 60 °C (60 s). All qPCR reactions were performed in triplicate with DNA template in the final replicate diluted 10-fold (up to 50-fold) to check for inhibition (44) and with additional replication as needed to repeat inhibited samples at the higher dilutions. The cycling protocol consisted of an initial denaturation at 95 °C for 10 minutes, followed by 40 cycles of 95 °C for 15 s and 60 °C for 60–90 s. The amplified PCR product was detected by monitoring the increase in fluorescence signal generated from the 6-carboxyfluorescein-labeled probe using a Realplex^2^ Mastercycler (Eppendorf). Data were analyzed using realplex software (Eppendorf) to determine cycle threshold (C_t_) values. A standard curve of serial 10-fold dilutions of genomic DNA (*V. vulnificus* strain YJ016) was run in triplicate along with the samples.

### Statistical treatment of data

Statistical analyses were conducted using JMP Pro 15 (SAS Institute, Inc.). Concentrations of *V. vulnificus* (CFU or *vvhA* gene copies mL^-1^), total bacteria, chl *a*, nutrients, PC, and PN were log transformed and rainfall and streamflow were cube-root transformed in order to normalize the data for multivariate analyses. For some analyses, sites were clustered into categories of “Upper canal” (Sites 1, 5–8) and “Lower canal” (Sites 10, 11, 13–15) based on whether they were landward or seaward of the sediment sill deposited at the mouth of the Mānoa-Palolo Stream. Comparison of means between two samples were conducted with t-tests assuming unequal variance or by the non-parametric Komlgorov-Smirnov Asymptotic Test for data that could not be readily normalized by transformation. Comparisons of means among three or more samples were conducted by ANOVA with a post-hoc Tukey-Kramer test of honestly significant difference. Factor analysis was conducted on *vvhA* and nutrient data using principal components with varimax rotation. For multiple linear regression, the data were split into two subsets (salinity < 12 or ≥12 ppt), because of the non-linearity in the relationship between *V. vulnificus* and salinity (21). Multiple linear regression models were also conducted on data covering the entire salinity range by either including a quadratic term for salinity (24, 45) or a derived variable ΔSal_opt_, which is the absolute value of difference between the sample salinity and an optimum salinity set as 12 ppt (46). Variables for constructing generalized regression models on each subset were selected using the Akaike Information Criterion by screening for the subsets that produced the best fit among all possible models. Among equivalent subsets in the “green zone” (AICc to AICc+4), either the subset with the best fit or with the fewest variables was selected as noted in the text.

Out of 243 total qPCR assays for *V. vulnificus* abundance, seventeen (ca. 7%) had issues that made them unreliable or unavailable (inhibition, below the reporting limit for the assay, or absence of data). In thirteen of these instances, abundances were instead inferred from blue colony counts on CHROMagar Vibrio medium (Supplemental Methods), based on the strong correlation (r = 0.8) between log-transformed concentrations of blue colony counts and vvhA gene copy numbers (Supplemental Fig. S1).

## RESULTS

### Variability of the habitat

Rainfall in Mānoa Valley, one of the major watersheds draining into the canal, varied from 0 to 15.8 cm in the 24-hour period preceding each sampling event. The average rainfall prior to samplings in the rainy season (Oct–Mar) was 3.7 cm, which was significantly higher (p = 0.0054) by an order of magnitude compared to the average in the dry season (Apr–Sep) of 0.27 cm (Supplemental Figure S2). Flow from the Mānoa-Palolo Stream varied from 0.4 to 2.6 m^3^ s^-1^ on sampling days and was strongly correlated with the prior 24-hr rainfall in Mānoa Valley (r = 0.87, n = 13, *p* = <0.0001; Supplemental Table S3). Both the canal-wide average salinity and temperature for the monthly samplings (n = 13) had significant negative correlations with prior 24-hr rainfall (r = -0.84, *p* = 0.0003 and r = -0.86, *p* = 0.0002, respectively).

Over the course of the 13-month study, measured surface water salinities in the Ala Wai Canal varied from 1 to 36 ppt (mean of 24 ppt) and temperatures from 19.2 to 31.8 °C (mean of 27 °C; Table 1). Salinity was highly variable throughout the study area reaching maxima of ≥ 29 ppt at every site and minima of ≤ 5 ppt at least once at each site except Site 15, which is the most seaward site in the harbor (minimum salinity of 11 ppt). As a consequence, there was no significant difference in average salinity among sites (ANOVA, *p* > 0.07). When samples were clustered by general location, average salinity in the upper and lower canal were not significantly different (*p* = .7818), but the combined stream mouth sites had significantly lower salinity on average than either the upper (p = .0083) or lower (p = .0016) canal sites.

**Table 1.** Variables measured on individual samples, the number of samples measured, and the geometric mean (geomean), mean, median, minimum (min) and maximum (max) values for each (reported to two significant digits).

| Variable | units | n | Geomean | Mean | Median | S.D. | Min | Max |
| --- | --- | --- | --- | --- | --- | --- | --- | --- |
| Temperature | °C | 242 | 27 | 27 | 27 | 2.6 | 19 | 32 |
| Salinity | ppt | 243 | 20 | 24 | 28 | 9.7 | 1 | 36 |
| Nitrate | μM | 209 | 18 | 37 | 17 | 48 | 0.02 | 260 |
| Nitrite | μM | 211 | 0.47 | 0.59 | 0.47 | 0.47 | 0.04 | 3.3 |
| Ammonia | μM | 207 | 5.5 | 6.7 | 5.7 | 4.3 | 0.94 | 22 |
| Phosphate | μM | 211 | 1 | 1.3 | 0.95 | 1.1 | 0.2 | 8.7 |
| Silica | μM | 211 | 110 | 140 | 120 | 92 | 11 | 490 |
| Particulate Carbon | μM | 199 | 130 | 250 | 120 | 650 | 20 | 7500 |
| Particulate Nitrogen | μM | 199 | 16 | 26 | 14 | 51 | 2.6 | 560 |
| Chlorophyll *a* | μg L<sup>-1</sup> | 194 | 7.6 | 19 | 7.3 | 43 | 0.4 | 500 |
| Total bacteria | 10<sup>9</sup> cells L<sup>-1</sup> | 219 | 4.2 | 4.8 | 4.7 | 2.3 | 0.47 | 11 |
| CaV blue<sup>1</sup> | CFU mL<sup>-1</sup> | 59 | 130 | 400 | 100 | 770 | 12 | 3904 |
| *vvhA* gene<sup>2</sup> | copies mL<sup>-1</sup> | 239 | 68 | 330 | 60 | 1200 | 3.4 | 14000 |

<sup>1</sup> culture-based blue colony forming units (CFU) when plating on CHROMagar Vibrio medium (CaV)
<sup>2</sup> qPCR-based estimates

All of the measured variables (Table 1) except silica and nitrite displayed overall significant positive or negative significant correlations with salinity (Supplemental Table S4), but the correlation coefficients were low in many cases because of non-linearity in the relationships (Fig. 2). Temperature displayed a significant, linear positive correlation with salinity (r = 0.65, n = 242, p < .0001). Correlation and regression analyses for all other variables vs. salinity are reported for log transformed data. Concentrations of chl *a* (range 0.4–512 µg L^-1^) showed a significant positive, linear (Fig. 2a) correlation with salinity (r = 0.49, n = 194, p < 0.0001). Concentrations of total bacteria (range 0.47 × 10^6^ to 11 × 10^6^ mL^-1^) also showed a significant positive correlation with salinity (r = 0.29, n = 219; p < 0.0001), but the relationship was non-linear (Fig. 2c). Particulate carbon (range 15–5,600 µM) had a non-linear relationship with salinity (Fig. 2d) that resulted in an overall weak but significant negative correlation (r = -0.25, n = 199, p = 0.0003).

**Fig. 2.**
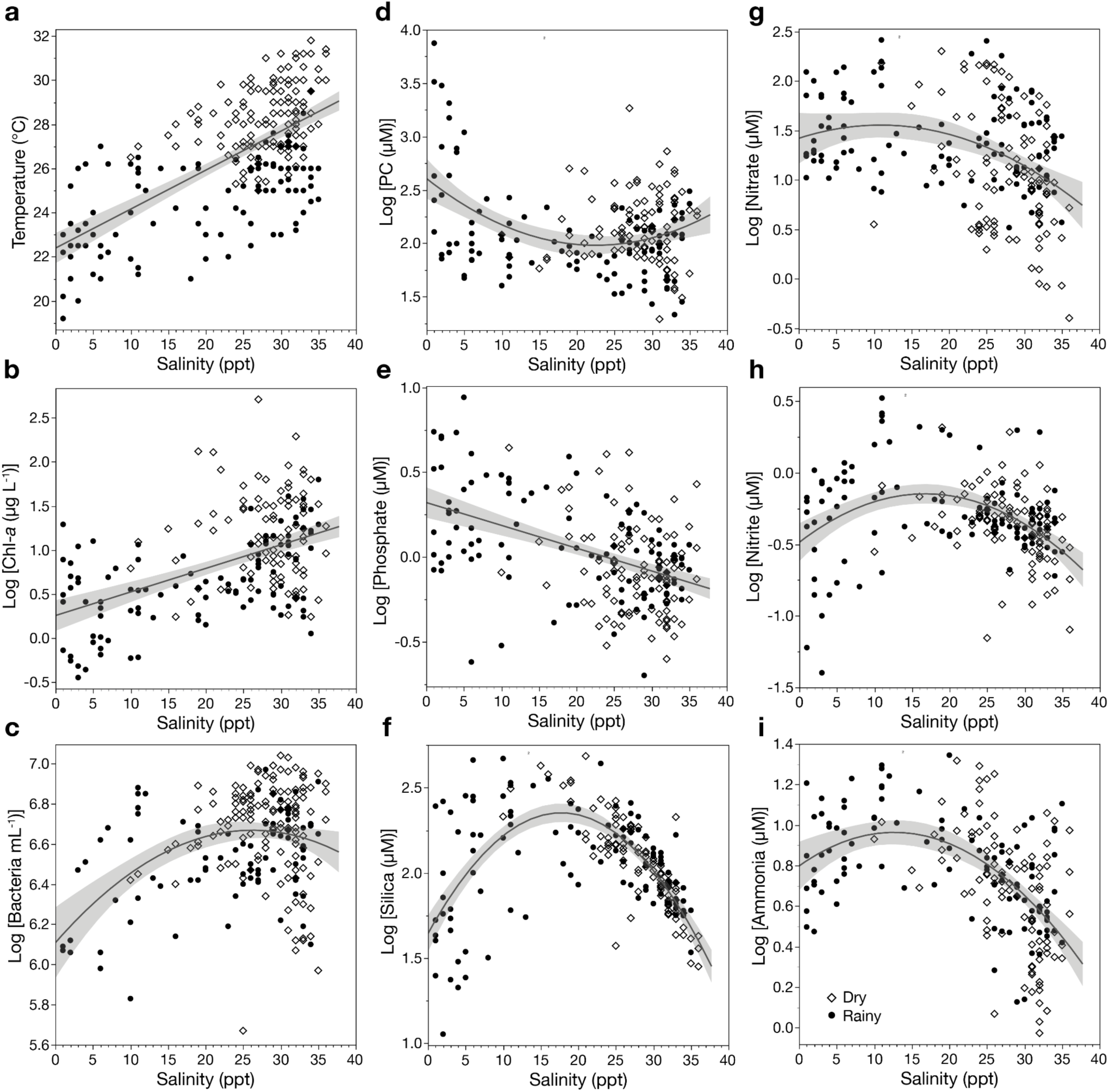
Variability in measured biological and chemical properties of samples as a function of salinity in samples from the rainy (solid circles) and dry (open diamonds) seasons. Regressions against salinity are shown for (**a**) temperature (r^2^ = 0.42), (**b**) Log chl *a* (r^2^ = 0.24), (**c**) log bacteria (r^2^ = 0.14), (**d**) log PC (r^2^ = 0.13), (**e**) log phosphate (r^2^ = 0.22), (**f**) log silica (r^2^ = 0.463), (**g**) log nitrate (r^2^ = 0.13), (**h**) log nitrite (r^2^ = 0.152), (**i**) log ammonia (r^2^ = 0.29). Regression lines and 95% confidence limits were fit using only first order terms unless addition of a quadratic term substantially improved r^2^ or reduced root mean square error. All fits were significant (p < .0001).

Of the dissolved inorganic nutrients, only phosphate (range 0.2–8.7 µM) had a linear relationship with salinity (Fig. 2e) and displayed a significant negative correlation (r = -0.46, n = 211, p < 0.0001). Concentrations of silica (11–490 µM), nitrate (0.02–260 µM), nitrite (0.04–3.3 µM), and ammonia (0.94–22 µM) all displayed significant, non-linear relationships with salinity (Fig. 2f-i), with highest values occurring at moderate salinities. Despite the non-linear relationships, there were significant negative correlations between salinity and either nitrate (r = -0.32, p < 0.0001) or ammonia (r = -0.44, p < 0.0001). Silica and nitrite, on the other hand, showed highly significant, non-linear relationships with salinity (Fig. 2f, h), that resulted in low and insignificant correlation coefficients.

When sites were clustered by location, most nutrients (nitrate, ammonia, phosphate, silica, but not nitrite), particulate carbon, chl *a*, and total bacteria were all significantly higher (p < 0.01) in the upper canal sites than the lower canal sites.

### Temporal and spatial variability of *V. vulnificus*

Concentrations of the *vvhA* gene (a proxy for *V. vulnificus* abundance) varied over four orders of magnitude in space and over time (Fig. 3) from 3 to 13,700 mL^-1^ with overall geometric mean concentration for all samplings of 68 mL^-1^ (n = 239; Table 1). Concentrations of *vvhA* at any given site were highly variable over time with values that were above average or below average occurring at some point at every location. Although spatial and temporal variability were low during the trihoral sampling over the course of one day in July, larger variations were seen on daily or longer time scales. The most dramatic variation is the change from above average to below average concentrations at every site in the span of 15 days (October 27 to November 11, 2008).

**Fig. 3.**
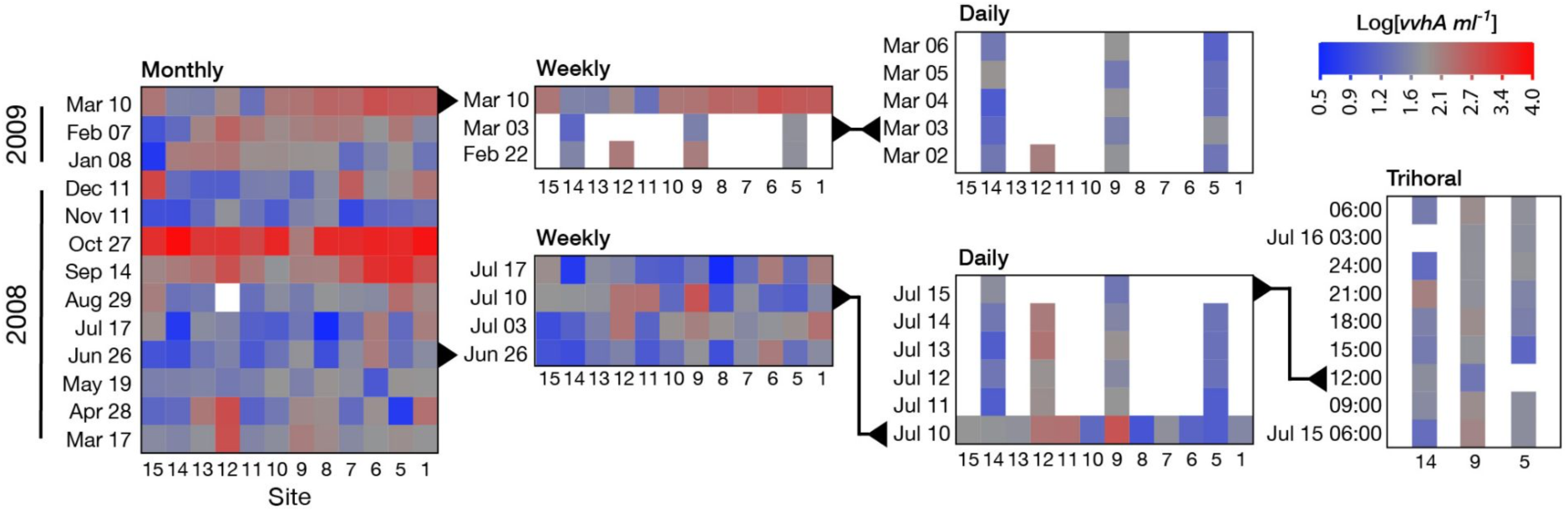
Heat maps illustrating spatial and temporal variability in *V. vulnificus*. Log *vvhA* concentrations are color coded at each station over time for monthly, weekly, daily and trihoral sampling events. The overall average log(*vvha*) from all samplings of 1.8 is shown in grey. Concentrations above average are in red and those below average in blue. The samplings on different time scales are nested and the events that are overlapping in the different graphs are indicated with black triangles and lines.

Despite the high variability, average log-transformed *vvhA* concentrations in the rainy season (1.98 ± 0.72) were significantly higher (unpaired t-test, *p* = .0013) than the dry season (1.72 ± 0.50; Supplemental Fig. S2). None of the individual sites had an annual average *vvhA* concentration that was significantly different from any of the others (ANOVA, post-hoc Tukey, *p* ≥ .63). However, excluding the stream mouth sites, the annual average concentration of log-transformed *vvhA* for the five sites in the upper canal (2.04 ± 0.74) was significantly higher (n = 65 at each site; unpaired t-test, *p* = .0110) than the annual average for five sites in the lower canal (1.75 ±0.70).

### Relationship of *V. vulnificus* to temperature and salinity

Log-transformed concentrations of *vvhA* displayed a weak, but significant, negative correlation (r = -0.174, *p* = .0071) with temperature (Fig 4a). However, partial correlation analysis indicates that the relationship between log[*vvhA*] and temperature is weakly positive, but significant (r = 0.258, *p* < .0001) when accounting for the effect of salinity and other variables. The relationship of log[*vvhA*] with salinity was non-linear with a peak around 12 ppt (Fig. 4b). Linear regression analysis with samples with salinity < 12 or ≥ 12 ppt showed that *vvhA* increased significantly (r^2^ = 0.315; F test, *p* = .0001) as a function of salinity over the lower range and decreased significantly (r^2^ = 0.492; F test *p* < .0001) over the higher range.

**Fig. 4.**
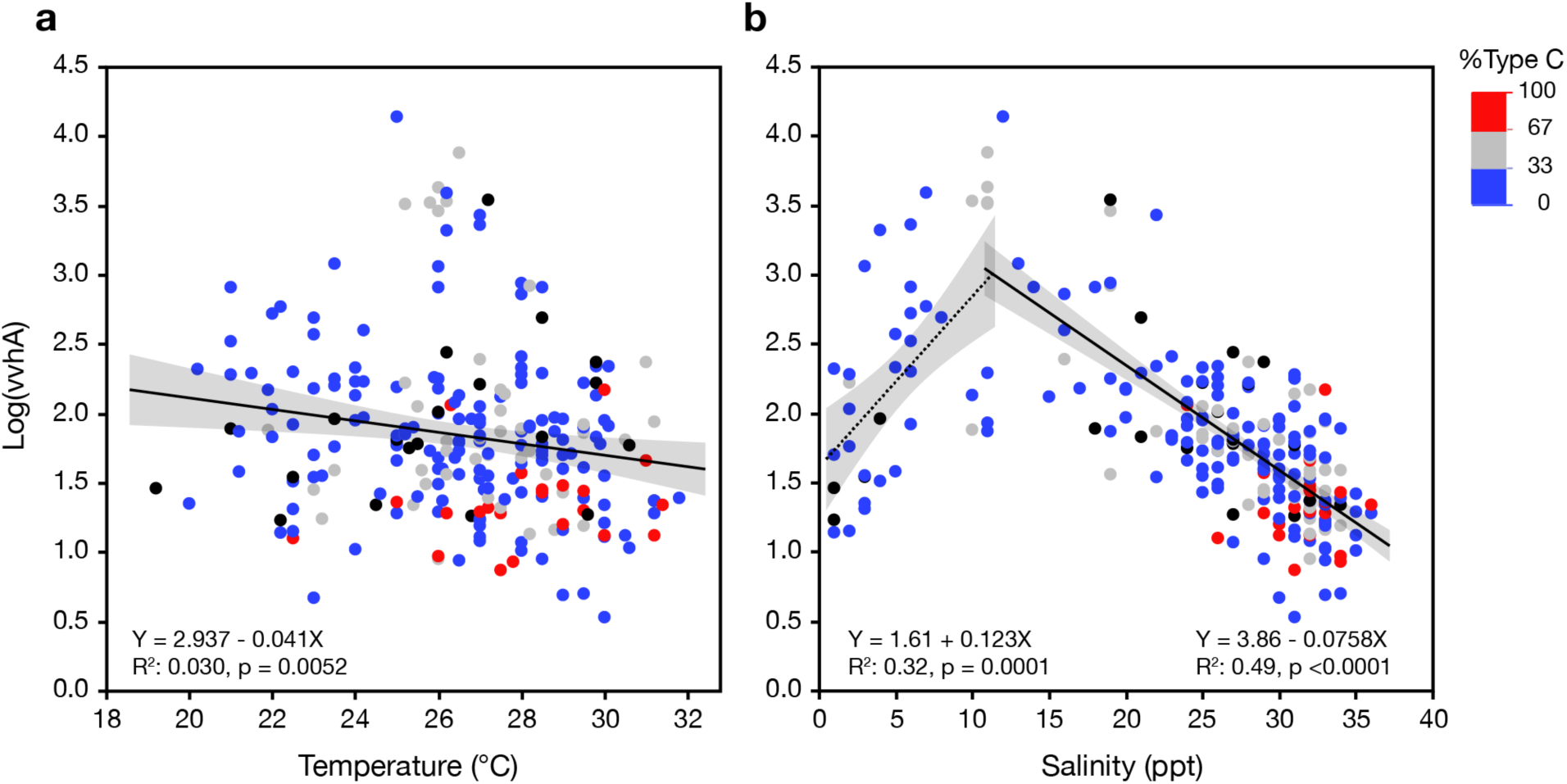
Concentration of *vvhA* as a function of **a**) temperature or **b**) salinity. The percentage of total *vvhA* that derives from “C-type” *V. vulnificus* was determined as the ratio of *vcgC* (C-type) and *vvhA* (total V. vulnificus) gene concentrations and is indicated by the color scale. Blue dots are samples dominated by E-type, red dots by C-type.

Concentrations of clinical, or C-type, *V. vulnificus* were usually lower than those of environmental, or E-type, and accounted for 26% of the total *V. vulnificus* on average across all samplings for which data were available (n = 219), indicating that communities were most often dominated by E-type. Both C-type and E-type *V. vulnificus* were most abundant at moderate salinities and declined as a function of salinity, but C-type declined at a lower rate. As a consequence, the contribution of C-type tended to increase as a function of salinity. Samples in which C-type accounted for >50% of the total (n = 39) were only observed in higher salinity waters (Fig. 4b) and the % C-type was significantly higher (Komolgorov-Smirnov; *p* = 0.0164) in higher salinity samples (≥ 25 ppt, n = 146) than samples having lower salinity (< 25 ppt, n = 73; Supplemental Fig. S3).

### Relationship between *V. vulnificus* and additional variables

To understand additional factors that may be important in controlling *V. vulnificus* in this habitat, factor analysis was conducted with *vvhA*, temperature, salinity, and nutrient data (Fig. 5a). Two factors had eigenvalues > 1. The strongest positive correlations (r ≥ 0.4) were between *vvhA* and silica or reduced nitrogen species, which were associated with Factor 1, and strong negative correlations (r ≤ - 0.4) were found between salinity and *vvhA*, ammonia, and phosphate along Factor 2. Plots of the factor loading values with points coded by rainfall and streamflow (Fig 5b) illustrate the relationship between these two related indicators of freshwater input and salinity along the Factor 2 axis. Coding the points by log *vvhA* concentration and silica concentration illustrates the association of these variables (along with reduced nitrogen species) with Factor 1. Overall the highest concentrations of vvhA occurred at moderate rainfall in the valley, but relatively low streamflow, and elevated concentrations of silica.

**Fig. 5.**
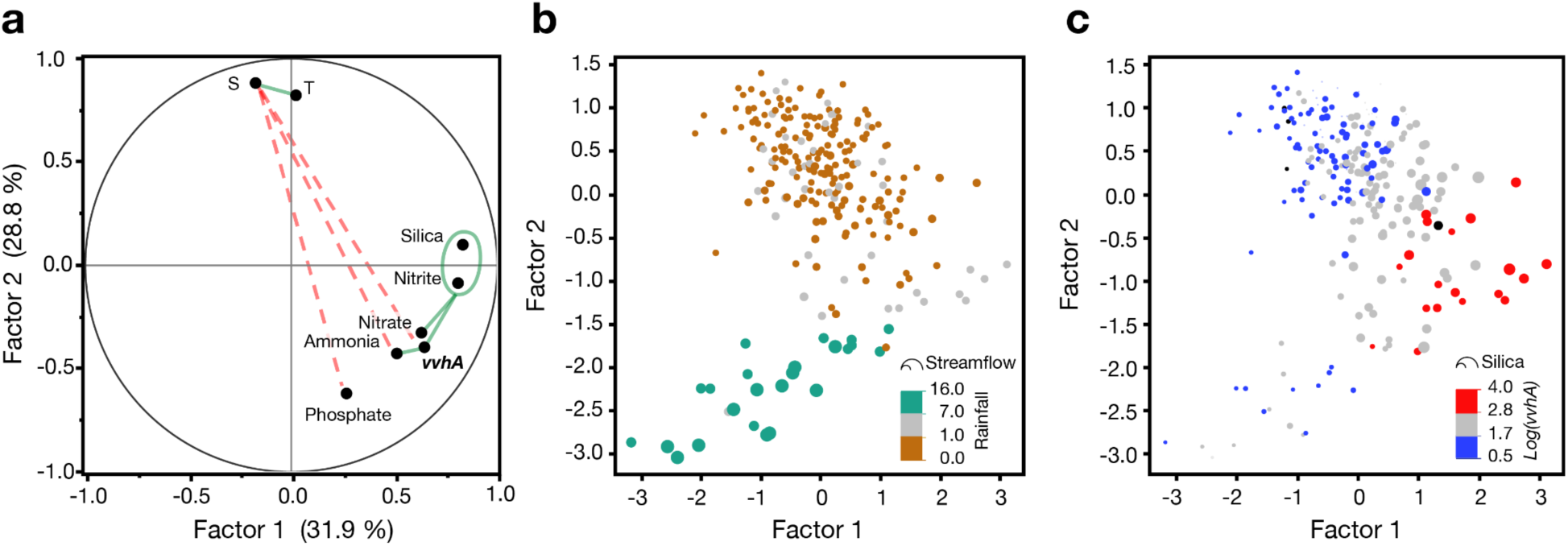
Factor Analysis for *vvhA*, temperature, salinity, and nutrients. (**a**) The factor loading plot for factors 1 and 2 (eigenvalues >1). Variables with a strong positive correlations (r ≥ 0.4) are connected by green solid lines and those with strong negative correlation (r ≤ 0.4) are connected by dashed red lines (**b**) Plot of the factor scores for the data with points colored by 24-hr antecedent rainfall in Mānoa Valley (in cm) and scaled in size so area is proporional to streamflow in the Mānoa-Palolo Stream. (**c**) Plot of the factor scores with data points colored according to log *vvhA* concentration and scaled in size so area is proportional to silica concentration.

Generalized regression models for predicting *vvhA* concentrations over the two different salinity ranges were constructed using the overall best subset (<12 ppt model) or the best subset having the minimum number of variables (≥ 12 ppt model). Only properties intrinsic to the individual samples were included in this analysis (i.e., tides, rainfall and streamflow were not considered). For samples with salinities < 12 ppt (n = 39 out of 41 samples, because of missing nutrient data) a subset of four (temperature, nitrite, silica, and PC) out of eight variables explained 75% of the observed variation with the equation:

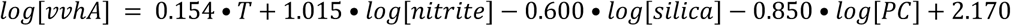

where *T* is temperature in °C, and nitrite, silica, and particulate carbon (PC) are in units of µM (model fit illustrated in Supplemental Fig S4a). For samples with salinities the ≥ 12 (n = 163 out of 198 possible samples because of missing nutrient data) a subset of just three (temperature, salinity and phosphate) out of seven variables explained 55% of the variability:

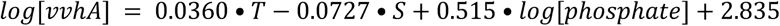

where *T* is temperature in °C, *S* is salinity is in units of ppt, and phosphate is in units of µM (model fit illustrated in Supplemental Fig. S4b). PC was removed prior to variable selection in the latter model, because initial analysis showed it offered no significant explanatory power at salinities >12 ppt, and missing data would have further restricted the samples included in the analysis. When predictions from the two models were combined, 66% of the variability in log(*vvhA*) over the entire salinity range was explained overall (Fig. 6).

**Fig. 6.**
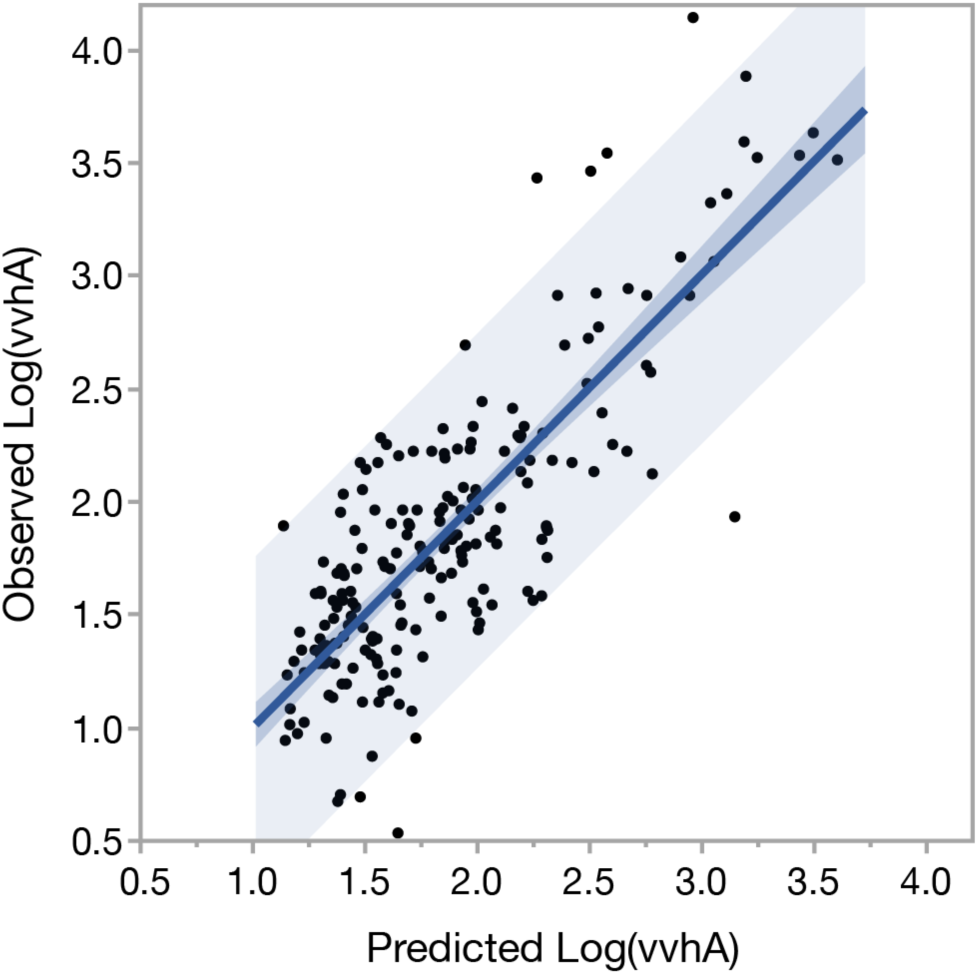
Observed vs. predicted values of log transformed *vvhA* gene copies per mL^-1^. Predicted values are combined from two separate models (one for samples < 12 ppt, one for ≥ 12 ppt). Predicted values are restricted to individual samples for which all of the predictor variables were measured within a given salinity range (n = 204 out of 239 in total). Darker and lighter shading illustrates the 95% confidence limits of the fit and and prediction, respectively. Combined, the models explain a significant amount of the variation in the observations: r^2^ = 0.661, RMSE = 0.37, F Test (1, 204) = 396.90, p < .0001.

Models in which either a quadratic term for salinity or the derived variable ∆ Sal_opt_ were included explained similar amounts of variability (r^2^ = 0.61 and 0.63, respectively; *p* ≤ .0001) using different sets of five variables (Supplemental Fig. S5), but were slightly outperformed by the combined models above.

### System-wide controls on *V. vulnificus*

To smooth out inter-station variability and focus on temporal variations in *vvhA*, canal-wide averages for the variables for each monthly sampling were also analyzed in relation to system-wide drivers of rainfall and streamflow (Fig. 7). In general, average rainfall, streamflow, phosphate, silica, and *vvhA* are all below average, and salinity above average, during most of the dry season with minimal variability. During the rainy season, periodic heavy rainfall resulted in high variability with excursions in all variables well above and below their overall averages.

**Fig. 7.**
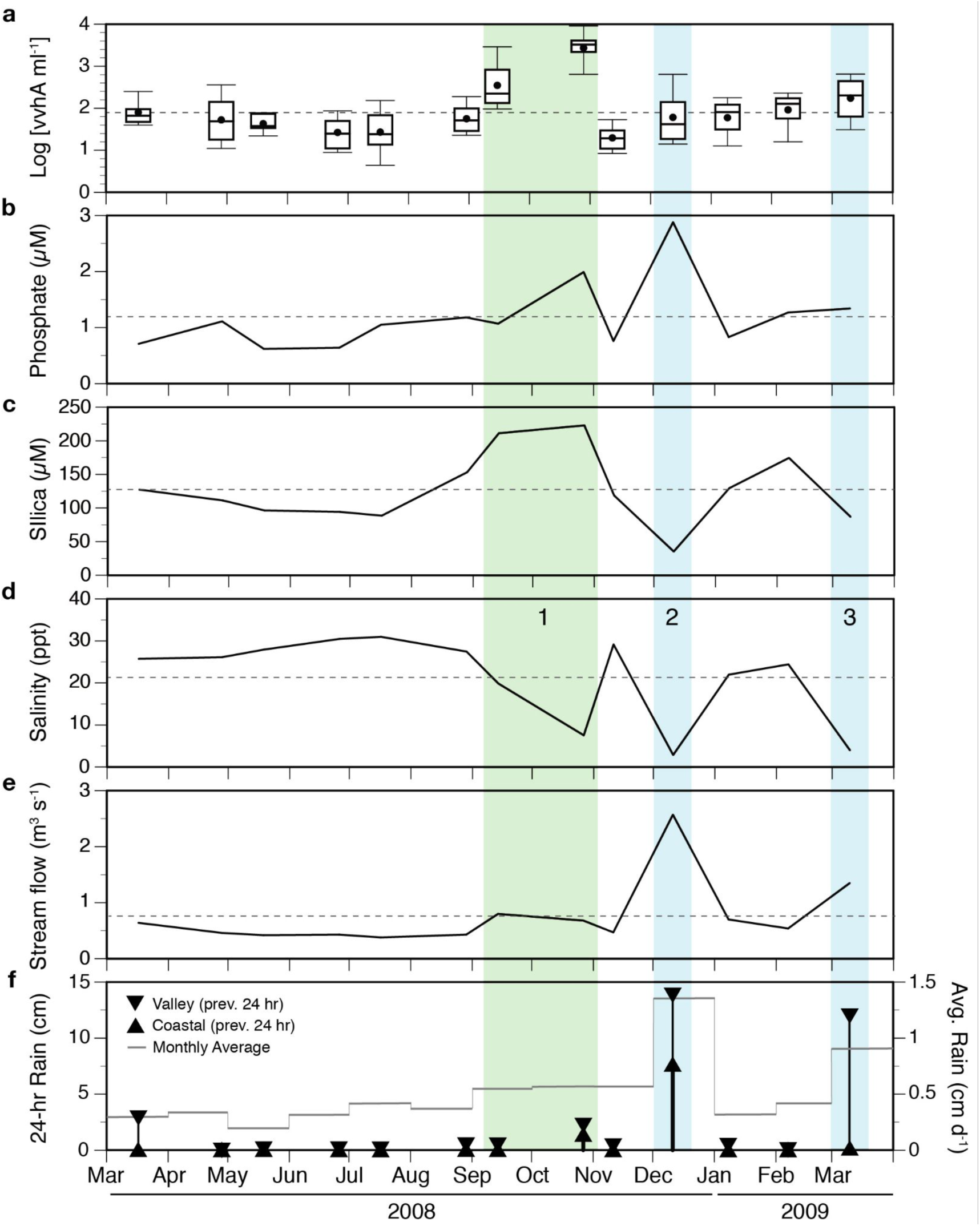
Time series of variables in or influencing the Ala Wai Canal system. Shown are **a**) variations of *vvhA* concentrations as box plots of all log transformed values measured at every site at each monthly sampling, canal-wide geometric means of **b**) phosphate, **c**) salinity, **d**) silica, as well as **e**) streamflow in the Mānoa-Palolo stream on the day of sampling, and **f**) rainfall in the 24-hr period preceding sampling as measured at the Honolulu coastal (upward triangles) and Mānoa Valley rain gauges (downward triangles). Daily rainfall average for the month is shown as the mean for both sites (grey line).

Three freshening events are evident from dips in the average salinity in the canal during the rainy season (Fig. 7). The first begins in September and peaks in October 2008 following increases in rainfall and streamflow. The average monthly rainfall increased from ≤ 0.75 cm d^-1^ in the preceding months to 1.0 cm d^-1^ in Sep– Oct, and the 24-hour antecedent rainfall for the October sampling was 2 cm (up from ≤ 0.5 cm in other samplings). Streamflow increased from 0.4 cm^3^ s^-1^ in July– August to 0.7–0.8 cm^3^ s^-1^ in Sep–Oct. Despite these relatively modest increases, canal-wide average salinity dropped to 8 ppt and the average silica concentrations in Sep–Oct reached their highest concentrations (223–244 µM). Phosphate displayed only a small local peak in average concentration (2 µM). Canal-wide average concentrations of *vvhA* reached a maximum during this event from 350 (range 67–3,500 gene copies mL^-1^ in September to an average of 2,700 (range 170 to 13,700) gene copies mL^-1^ in October. The average concentration in October was significantly higher than at any other monthly sampling (ANOVA, post-hoc Tukey, *p* ≤ .0005). At the subsequent sampling 15 days later (November), rainfall had stopped, streamflow, phosphate and silica had declined, average salinity had increased to 29 ppt and *vvhA* was at the lowest average concentration of the study with an average of 20 (range 7–63) gene copies mL^-1^ across all sites.

A second, more pronounced drop in salinity occurred in December 2008 in response to heavy rainfall recorded at both the coastal and Mānoa valley rain gauges, resulting in the highest recorded streamflow (2.6 m^3^ s^-1^), minima in salinity (3 ppt) and silica (34 µM), and the highest average phosphate concentration (2.9 µM). In contrast to the previous freshening event, *vvhA* was not significantly elevated (61 gene copies mL^-1^) and was near the overall study average.

A third freshening event occurred at the time of the last sampling in March 2009 as a result of heavy rainfall in Mānoa Valley, but not at the coast. Streamflow (1.3 m^3^ s^-1^) was above average and intermediate between the first and second events, and salinity was again significantly reduced (4 ppt). The effects on phosphate (1.3 µM) and silica (87 µM) were modest, with phosphate being just above the long-term average and silica just below. The mean concentration of *vvhA* reached its third highest level at this time reaching 175 (range 22–811) gene copies mL^-1^ after steadily increasing each month from the lowest value in November.

Multiple linear regression was used to determine which subset of variables best predicted canal-wide average log(*vvhA*) concentrations. The model resulting from the best subset out of all combinations of twelve possible variables was:

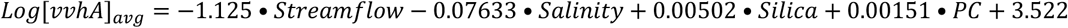

where streamflow is in units of m^3^ s^-1^, and salinity, silica, and particulate carbon (PC) are in units of µM. All variables are the geometric means for all sites in the canal for each monthly sampling (n= 13). Linear regression of the observed vs. predicted *vvhA* suggests that 97% of the canal-wide average variation in vvhA could be explained with the selected variables (Fig. 8a).

**Fig. 8.**
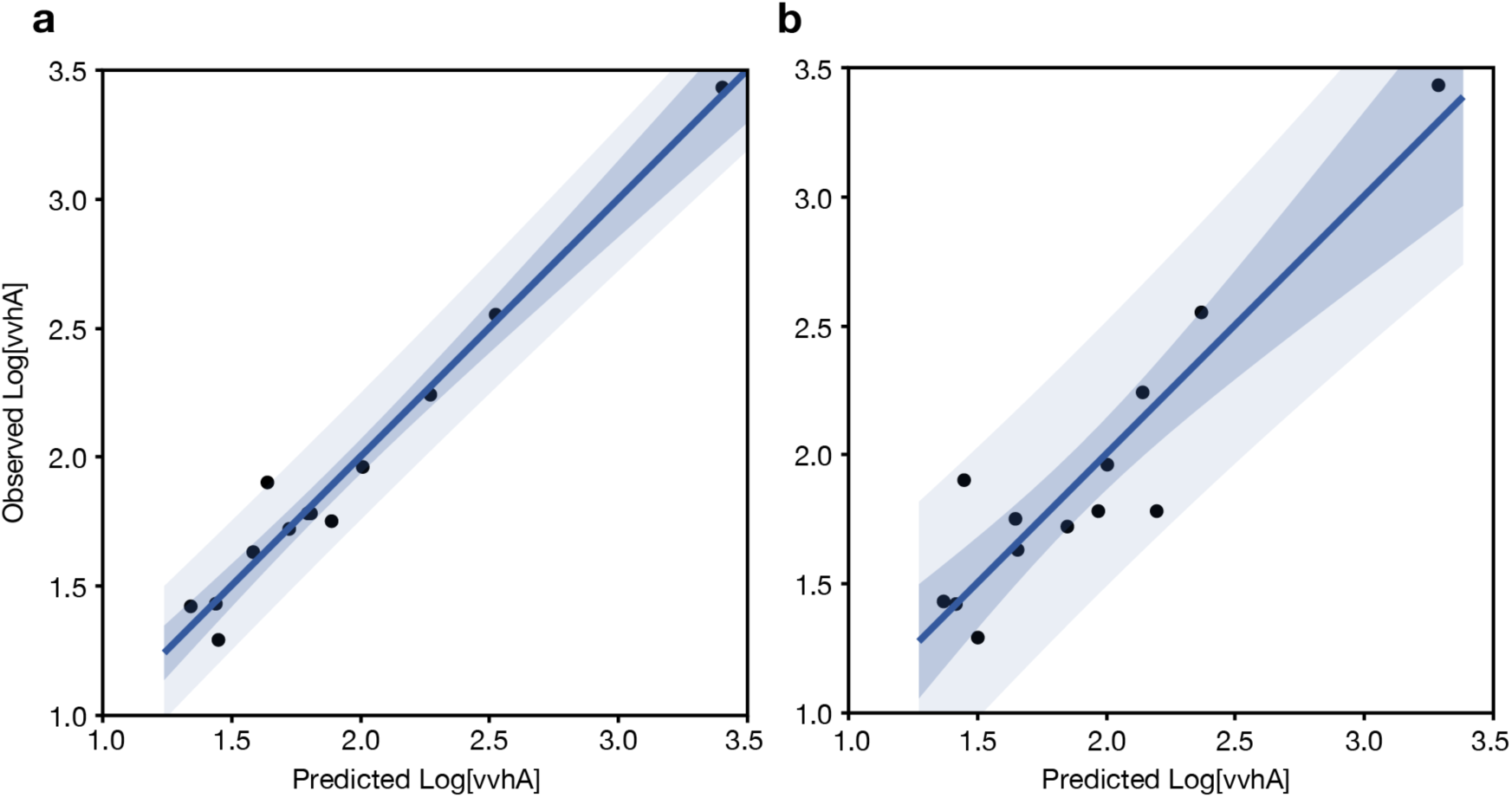
Observed vs predicted canal-wide average of log-transformed *vvhA* concentrations. Predictions are derived from **a**) the best subset of variables (salinity, silica, streamflow, particulate carbon) from generalized regression model (r^2^ = 0.97; RMSE = 0.11; F test p < .0001) or **b**) a restricted subset of two variables (rainfall and salinity) that are easily measured autonomously (r^2^ = 0.86; RMSE = 0.22; F test p <.0001). Darker and lighter shading illustrates the 95% confidence limits of the fit and and prediction, respectively.

A second simpler model using a minimum of readily measurable variables (salinity and rainfall) was also constructed:

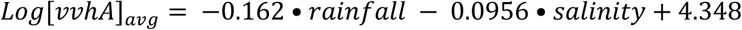

where rainfall is average rainfall in cm for the prior 24 hours at the Mānoa Valley gauge, and salinity is canal-wide average salinity in ppt. This simpler model explained 83% of the variability in average log-transformed concentrations of *vvhA* (Fig. 8b).

## DISCUSSION

### Temporal and spatial variability of *V. vulnificus*

*V. vulnificus*, as inferred from *vvhA* gene, was consistently detected throughout the year in the Ala Wai Canal and Harbor system, but varied dramatically over space and time. Sampling on different temporal scales showed minimal variation in *V. vulnificus* within a day, but dramatic and stochastic variations on longer time scales and among sites. This suggests that factors with regular intra-day variations (e.g. tides, or daily changes in temperature and primary productivity driven by insolation) had relatively little influence on concentrations of *V. vulnificus*. The largest absolute change in the canal-wide average *vvhA* concentrations seen over the entire study occurred in a span of 2 weeks. The observation that *V. vulnificus* concentrations were higher on average in the rainy vs. dry season, yet the lowest average concentration recorded in the study also occurred in the rainy season within weeks of the highest abundances, suggests that freshwater input, which occurs stochastically, but with an underlying strong seasonal component, is the most significant contributor to variability in *V. vulnificus* abundance in this environment.

The results support an earlier hypothesis (21) that in tropical and some subtropical climates, where the temperature range is narrow and persistently warm, salinity is a stronger determinant than temperature of *V. vulnificus* abundance. This is consistent with the seasonal variation in *V. vulnificus* in oysters in India, which is not related to temperature, but by summer monsoonal rains lowering salinity (47). In Hawaiʻ i, with its rainy season in winter months, there is a tendency toward a seasonal cycle in *V. vulnificus* abundance that is inverted from the pronounced temperature-driven cycle found at higher latitudes and the monsoon-driven cycle in India.

### Variable sources and influence of freshwater inputs

The two major sources of freshwater to the Ala Wai canal are surface runoff (primarily point source from streams and storm drains) and groundwater seeps. Compared to surface runoff, groundwater in Hawaiʻ i tends to be enriched in silica as a result of prolonged water-rock interactions (48) and depleted in phosphate as a result of interactions with lateritic soils containing high concentrations of iron and aluminum oxyhydroxides (48, 49). These differences, along with information on rainfall and streamflow, are helpful in identifying the primary source of the freshwater entering the canal. In the factor analysis, Factor 1 may be interpreted as a latent variable for groundwater (high loading for silica, but low for phosphate), and Factor 2 as a latent variable for (negative) surface runoff (high, but opposing, loading of salinity and phosphate). Plots of the loading scores reinforce the observation that *V. vulnificus* tended to be highest at moderate salinities and suggest that groundwater was a relatively more important source of freshwater input under those conditions (low streamflow, but elevated silica). When rainfall was highest, surface runoff contributed more to freshwater input (highest streamflows with high phosphate, low silica) and was associated with lower concentrations of *vvhA*.

This variable relationship between *vvhA* and freshwater source was also discernible in the temporal changes in variables when averaged across all canal sites. Of the three major freshening events, the first, with relatively high silica and low phosphate, suggests a significant contribution from groundwater. This is consistent with the observation of significant freshening, despite only modest increases in stream flow compared to the summer months. This presumed increase in groundwater input appears to have been driven by a moderate increase in monthly average rainfall in both September and October, coupled with a modest increase in average rainfall during the 24 hours preceding sampling that was greater at higher elevations in the watershed, than locally.

The second freshening event, with a high concentration of phosphate but low silica, appears to be dominated by surface runoff, resulted from a Kona storm on the south shore of Oahu (38). A Kona storm is a rain event that deviates from the normal northeasterly trade-wind driven patterns that govern Hawaii’s weather, and occurs when southwestern Kona winds bring heavy rains to the southern shore of Oahu. This storm resulted in unusually high rainfall, both higher in the watershed and locally, in the 24 hours prior to sampling.

The third freshening event on March 10, 2009 appears to have a source signature that is intermediate to the two prior events in terms of stream flow and silica. This is consistent with an average rainfall in the preceding 24 hours in the watershed that was high enough to increase downstream runoff and groundwater discharge into the canal (as in the previous event), but with limited local precipitation that, unlike the previous event, did not contribute appreciably to surface runoff.

The concentrations of *vvhA* during these three events suggest that the magnitude, if not the sources, of the freshwater input to the canal has a large influence on *V. vulnificus* abundances. The mixing of freshwater with the seawater in the canal is expected to have competing influences on *V. vulnificus*, because it simultaneously alters temperature, salinity, and residence time. At sustained, moderate levels of freshwater input (such as that from ground water intrusion driven by moderate rainfall higher in the watershed), both the temperature drop and decrease in residence time are relatively small, but the freshening is sufficient to result in salinities that are optimal for *V. vulnificus*, thus explaining the unusually high abundance of *V. vulnificus* in September and October 2008. During unusually intense storms, especially with high rainfall lower in the watershed (December 2008), the very high levels of surface runoff appear to suppress abundances of *V. vulnificus* in the canal. This is likely a result of the simultaneous reduction in growth rate (caused by decreases in both temperature and salinity to below optimum) and reduced residence time of water in the canal. Gonzalez {Gonzalez:1971to} observed an inverse relationship between streamflow and residence time of runoff waters in the Ala Wai Canal.

Although intense storms can temporarily suppress the canal-wide average concentrations of *V. vulnificus* in the canal/harbor system, the actual changes are site-specific. We observed, for example, that during the December 2008 storm, *V. vulnificus* abundance, despite a lower canal-wide average, was higher than average at Site 15, the most seaward site located in the Ala Wai Boat Harbor. In this location, salinity was temporarily reduced to 13 (in the optimal range for *V. vulnificus*) compared to the typical average salinity for this site of ≥ 30 (38). Salinity remained below the average in the harbor for 16 hours following the cessation of rainfall. This suggests that the sites posing the highest risk of infection by *V. vulnificus* will vary depending on the rainfall patterns and can even include the harbor which usually had some of lowest concentrations. This condition-dependent elevated risk in the harbor is consistent with the unfortunate incident of infection and death of an individual who had open wounds exposed to harbor water following a long period of intense rainfall (50).

### Patterns of *V. vulnificus* strain abundance

C-type *V. vulnificus* are the strains most frequently associated with infections in humans (9), but are often less abundant than E-type in environmental samples (9, 51). This appeared to be the case in our study site with C-type *V. vulnificus* accounting for an estimated 25% on average. The percentage was highly variable, however, and our observation that the C-type *V. vulnificus* tended to make up a higher percentage of the total at higher salinities (despite declining in absolute abundance) is consistent with some previous observations. Williams et al. (2017), for example, observed a negative influence of salinity on the abundance of E-type and C-type strains, but the effect was greater for E-type (51). Lin and Schwartz (2003) observed that when temperature decreased and salinity increased, in situ abundance of 16S rRNA A-type strains (analogous to E-type) decreased while B-type (analogous to C-type) increased and temporarily became the dominant genotype (52). Other studies in high salinity (> 32 ppt) coastal waters have found that either a majority (7) or all (53) of the isolates obtained were of B-type (C-Type). These observations support the contention that these different genotypes reflect distinct ecotypes, with the C-type having greater stress tolerance (10).

Multiple linear regression analysis was used to model *V. vulnificus* abundance using a reduced number of variables. Although these variables explained a significant percentage of the variability in *V. vulnificus* abundance, a great deal of sample-to-sample variability remains unexplained, which is not uncommon (21, 26). Predicting system-wide average concentrations of *V. vulnificus*, on the other hand, was much more successful. A model with the best subset of four variables explained 97% of the variability, and much simpler model relying on only two readily obtainable measurements (rainfall and salinity), still accounted for much of the variability and might prove more useful in practice for predicting relative risk from *V. vulnificus* of exposure to waters of the canal and harbor.

The high level of predictability for system-wide average *V. vulnificus* is similar to that achieved using logistic regression to predict *vvhA* as a binary response variable either as presence vs. absence (46) or low vs. high abundance (54). Improvements in the prediction of *V. vulnificus* at higher resolution may be realized by combining biological population models for *V. vulnificus* with physical models of coastal circulation (54). In the meantime, the results from this study provide a detailed description of the ecology of *V. vulnificus* in tropical estuarine waters of Hawaiʻ i. The results are a useful first step toward predicting and, ultimately taking steps to mitigate, the incidence of *V. vulnificus* infections.

## ACKNOWLEDGMENTS

We are grateful to G. Walker and B. Marchant for assistance with sample collection and A. Culley for support and advice. We thank Hawaiʻ i Ocean Time-series program staff for support with processing PC/PN samples and R. Briggs for advice on chemical measurements. This work was supported by grants from Hawaiʻ i Sea Grant (2009, 2012) and the National Science Foundation (OCE05-54768, OCE08-26650) and NOAA Ocean Observing (NA07NOS4730207).

